# Raman mineral-to-matrix ratios correlate with weight percentage mineral-to-matrix ratio determined by in-SEM Raman imaging of bone tissue

**DOI:** 10.1101/2024.05.30.596667

**Authors:** Guillaume Mabilleau, Dale Boorman, Jorge Diniz

**Affiliations:** Univ Angers, Nantes Université, ONIRIS, Inserm, RMeS, UMR 1229, F-49000, Angers, France; CHU Angers, Bon pathology unit, 49933 Angers, France; Renishaw plc, New Mills, Wotton-under-Edge, Gloucestershire, GL12 8JR, UK

**Author notes:** Correspondence to: Dr Guillaume Mabilleau, Inserm UMR_S 1229 RMeS – REGOS Team, Institut de Biologie en Santé, Université d’Angers, 4 rue Larrey, F-49933 Angers, France.

**Keywords:** Bone material properties, In-SEM Raman imaging, mineral-to-matrix, quantitative backscattered electron imaging

## Abstract

Raman imaging combined with scanning electron microscopy (SEM) is a powerful technique that allows for topographical, chemical and structural correlative multi-scale imaging. It provides the perfect tool to determine which of the Raman mineral-to-matrix ratios represent the best parameter to accurately measure the degree of mineralization of the bone matrix using quantitative backscattered electron imaging (qBEI) as the reference methodology. Indeed, previous studies evidenced that the v_2_PO_4_ and v_4_PO_4_ vibrational modes were less sensitive to laser polarization than the v_1_PO_4_. However, using the v_2_PO_4_ or v_4_PO_4_ requires a longer acquisition time or lower spectral resolution. In the present study, we evaluated the correlation between mineral-to-matrix ratios computed from v_1_PO_4_ and v_2_PO_4_ in a human bone sample retrieved from orthopaedic surgery during hip replacement and wt% mineral / wt% organic matrix obtained from qBEI using the inLux SEM Raman interface. We reported here that all mineral-to-matrix ratios were significantly linearly correlated with wt% mineral / wt% organic matrix and that v_1_PO_4_/CH_2_ exhibited the strongest correlation coefficient (r=0.880). This study suggests that the v_1_PO_4_ is still a valid Raman peak to estimate the mineral-to-matrix ratio in bone samples and can be used to diagnose bone fragility disorders.

## 1. INTRODUCTION

Bone is a living material made of an organic phase composed mainly of type I collagen fibres and interfibrillar spaces filled in with non-collagenous proteins, poorly crystalline hydroxyapatite crystals and water ^1^. The major roles of bone tissue are to act as a reservoir of calcium and phosphate but also to adapt continuously to mechanical stress by modulating bone mass, microarchitecture and bone material properties ^2^. The latter has attracted attention as an important contributor of bone strength at the organ and tissue levels. At the organ level, the mineral-to-matrix ratio of the bone matrix has been positively correlated to ultimate stress, bending modulus and stiffness, and it has been inversely correlated to post-yield displacement and toughness in rodents and human bones ^3-7^. At the tissue level, the mineral-to-matrix ratio has also been positively correlated to indentation modulus, hardness and plastic index in rodent and sheep bones ^5,8-12^. Several indicators of the degree of mineralization have been proposed based on x-ray absorptiometry, scanning electron microscopy (SEM) and vibrational spectroscopies. The latter are sensitive to chemical and structural properties of the specimen, providing qualitative and quantitative information about sample composition. Raman spectroscopy is a well-suited methodology for the analysis of bone material and has many advantages over other analytical techniques: it is non-contact and non-destructive, highly sensitive to molecular and structural properties (including many functional groups), requires little sample preparation, enables label-free biochemical detection, and is compatible with aqueous matrices since it has negligible interference from water. In Raman spectroscopy, several indices of mineral-to-matrix ratio have been proposed based on the v_1_, v_2_ and v_4_ vibrational modes of the PO_4_ group and weighted by vibrational modes assigned to the organic phase (Amide III, Amide I, amino acid side chains) ^13^. However, since bone is a birefringent material, the relative intensity of the mineral and organic peaks in Raman spectra depends not only on composition but also on the orientation of collagen fibrils relative to the polarization axis of the Raman instrument ^13^. This polarization bias can be minimized by using low numerical aperture objective (NA < 0.4) and bone cross-section ^13,14^. Two peak ratios have been reported as being less sensitive to polarization bias, namely the v_2_PO_4_/Amide III and the v_1_PO_4_/Proline because these peaks are in-phase _15,16_. Although the v_2_PO_4_/Amide III peak ratio has previously been correlated with the calcium content measured by quantitative backscattered electron imaging (qBEI), the mineral-to-matrix ratio using the v_1_PO_4_ has been less well-studied.

Recent developments in Raman instrumentation have allowed for a Raman interface to be fitted directly in a scanning electron microscope chamber allowing for simultaneous and accurate correlation of information from both modalities. The high resolution and diversity of signals for electron imaging (secondary, backscattered and Auger electrons) can greatly complement the chemical and structural information from the diffraction-limited Raman instruments, and vice-versa. Notably, the complementary potential of this multimodal approach is enhanced when Raman images are obtained at high spatial resolution, since chemical features in the specimen can be more accurately detected. This is of particular interest in the correlation between the degree of mineralization measured by electron microscopy, as done in qBEI, and the mineral-to-matrix ratio measured by Raman spectroscopy.

The aim of the present study was to assess the calcium content of a femoral neck cross-section by qBEI and find the best correlation to the mineral-to-matrix ratio used to describe the degree of mineralization, evaluated using Raman spectroscopy. To achieve this, a novel approach was tested using an innovative system for in-SEM Raman spectroscopy.

## 2. MATERIALS AND METHODS

### 2.1. Bone sample preparation

The bone sample was collected from a 64-year-old man suffering with type 2 diabetes mellitus and admitted to the orthopaedic surgery department at Angers University Hospital for a femoral neck fracture. The proximal end of the femur was retrieved during total hip replacement surgery for routine bone pathology analysis. The patient gave their written informed consent for the use of this sample for research purposes. The bone sample was fixed in alcoholic formalin for 24 h prior to dehydration with acetone and delipidation with xylene. The specimen was then embedded undecalcified in polymethylmethacrylate at 4°C as previously reported ^17^. After hardening, a thick cross-section was made with a diamond saw and the specimen was ground with sandpaper with increasing grit-number followed by surface polishing with diamond paste (grain size down to ∼1 µm). The surface was then carbon coated using a Leica ACE600 device (Leica ACE600, Nanterre, France). The thickness of the carbon layer was measured with a quartz crystal during coating and estimated at ∼10 nm.

### 2.2 Quantitative backscattered electron imaging (qBEI)

Quantitative backscattered electron imaging was performed in high vacuum mode at ∼2.10^−4^ Pa in order to determine the bone mineral density distribution (BMDD) as previously reported ^18^. Briefly, the bone specimen was imaged with a scanning electron microscope (EVO LS10, Carl Zeiss Ltd, Nanterre, France) equipped with a five quadrant semi-conductor backscattered electron detector. The microscope was operated at 20 keV with a probe current of 250 pA and a working distance of 15 mm. The probe current was monitored in the chamber with a Faraday cup and a picoamperemeter. The backscattered signal was calibrated using pure carbon (Z = 6, mean grey level = 25), pure aluminium (Z = 13, mean grey level = 225) and pure silicon (Z = 14, mean grey level = 253) standards (Micro-analysis Consultants Ltd, St Ives, UK). For these contrast/brightness settings, the BSE grey level histogram was converted into weight percentage of calcium as described in Roschger et al. ^19^. Changes in brightness and contrast due to instrument instabilities were checked by monitoring the current probe and imaging the reference material (C, Al and Si) every 15 min. The region of interest (ROI) was located in the haversian bone. The ROI was imaged at a 50× nominal magnification, corresponding to a pixel size of 2.25 μm and a dwelling time of 100 µs/pixel. The grey level distribution of each image was analysed with a lab-made routine using Matlab 2023b (The Mathworks, Natick, MA). Five variables were obtained from the bone mineral density distribution: Ca_peak_ as the most frequently observed calcium concentration, Ca_mean_ as the average calcium concentration, Ca_width_ as the width of the histogram at half maximum of the peak, Ca_low_ as percentage of bone area which is mineralized less than 15.65 wt% Ca, and Ca_high_ as the percentage of bone which is mineralized above 24.99 wt% Ca. Thresholds for Ca_low_ and Ca_high_ were determined based on the BMDD data obtained on bone biopsies of a reference population of healthy volunteers available at the bone pathology unit at Angers University Hospital ^20^.

The BMDD of the bone specimen was further reinvestigated in a scanning electron microscope (Jeol JSM 6060LV, Jeol, Welwyn Garden City, UK) operated in low vacuum mode at 50 Pa. The microscope settings were as follows: accelerating voltage: 20 kV, probe current: ∼1 nA, working distance: 17 mm, dwelling time: 130 µs/pixel. The bone sample was imaged at a 100x nominal magnification, corresponding to a pixel size of ∼1 µm/pixel. The backscattered signal was calibrated using pure carbon, pure aluminium and pure silicon as described above. Given that qBEI was now conducted in low vacuum conditions, we opted to eliminate the carbon layer coating on the specimen by surface grinding and polishing with sandpaper and diamond paste (grain size down to ∼1 µm), as it was no longer necessary. Additionally, this aided in heat dissipation during Raman acquisition and hence avoided sample degradation. The BMDD parameters were computed using the same Matlab routine as for images obtained at the University of Angers.

As Raman spectroscopy reports the amount of mineral weighted by the amount of organic matrix, we computed the weight percent mineral (wt% mineral) from the qBEi image using the approximation of pure hydroxyapatite of known composition Ca/P = 1.67. As such, wt% mineral is estimated as 2.51x wt% Ca ^21^. We also made the assumption that the pMMA contribution in mineralized bone and the contribution of Ca and PO_4_ outside the mineral phase are negligible. As such the amount of organic matrix can be derived from qBEi image as 100 – wt% mineral. A full description of the method is provided in Roschger et al. ^22^. Wt% mineral over 100-wt% mineral map was computed from qBEI image.

### 2.3. Raman spectroscopy

Raman spectroscopy was conducted inside the SEM chamber (Jeol) with the Renishaw inLux™ SEM Raman interface (Renishaw Plc, Wotton-under-Edge, UK). This interface allowed for in-SEM Raman analysis of the same ROI without moving the sample. The inLux interface (Figure 1) was coupled to the side port of the SEM microscope (Jeol) and used a motorised probe to deliver and collect light from the sample, which also enabled *in-situ* Raman mapping. The Raman probe could be inserted to be aligned with the SEM electron beam for simultaneous Raman and secondary electron (SE) imaging on the same ROI on the sample. The Raman probe could be retracted to allow BEI detection, while the sample remained in place. Laser delivery and Raman light collection was allowed *via* a parabolic mirror in the Raman probe tip; in addition, it could also collect optical images using optical lenses, with a magnification similar to a 50× long-working distance objective and a numerical aperture of approximately 0.5. Simultaneous SE and Raman imaging was possible *via* a pinhole in the parabolic mirror, at both high and low vacuum conditions, which were used to confirm co-localisation between the different modalities.

**Figure 1:**
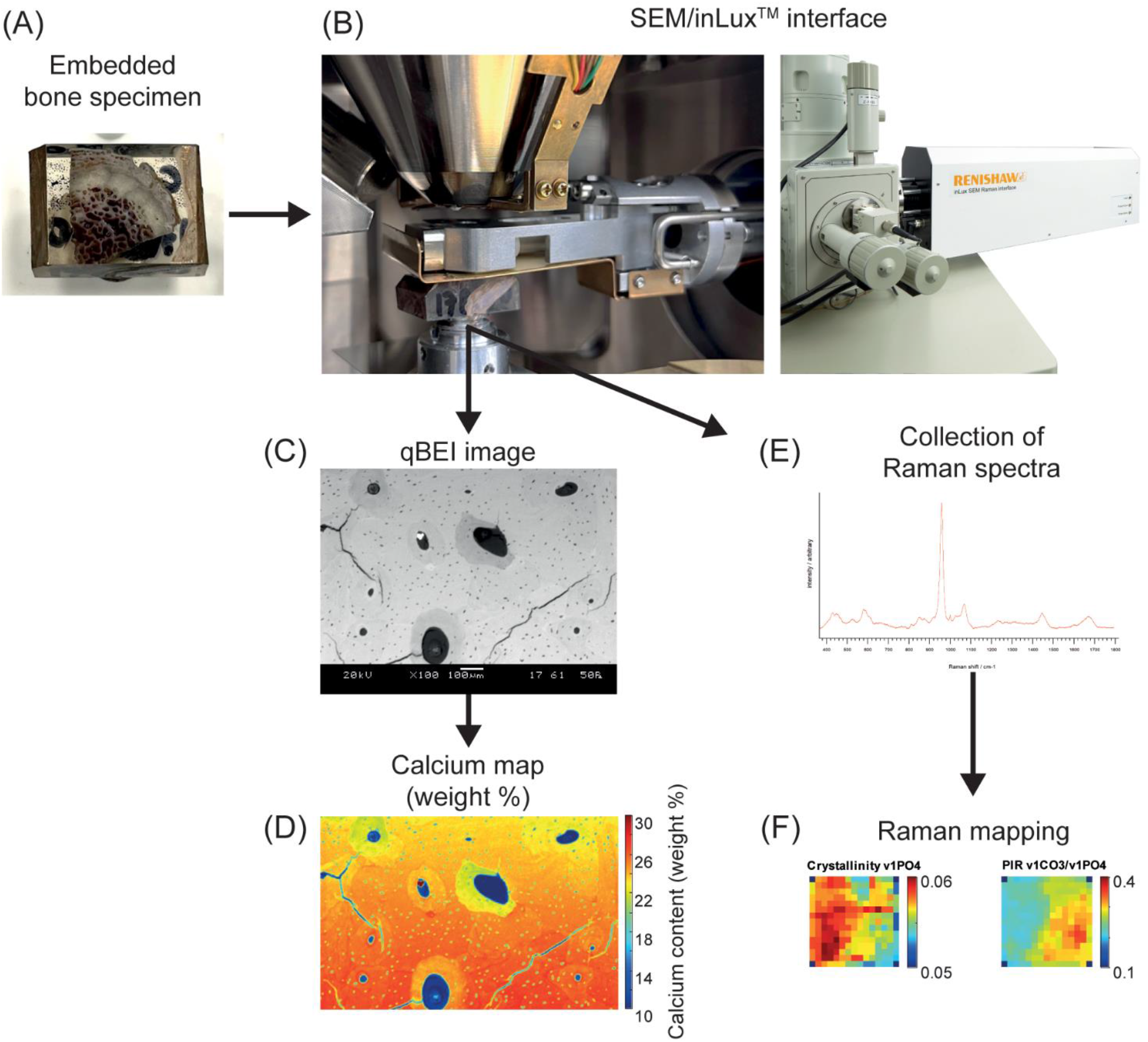
Schematic of the scanning electron microscope (SEM) / Raman inLux – Virsa system. (A) the bone sample was embedded in polymethymethacrylate and surface polished. (B) The bone specimen was introduced in the SEM chamber under the Raman probe and the electron beam pole piece. The probe was fully inserted in the SEM chamber and was aligned with the electron beam. It could be retracted for backscattered electron imaging, eliminating the need to move the sample between analysis for accurate co- localised work. The inLux interface was attached to a side port of the SEM chamber and coupled to the Virsa Raman spectrometer (not shown) via fibre optic cables. (C) A qBEI image was acquired and (D) converted into a calcium map. (E) Without moving the SEM stage, Raman spectra were acquired through the Raman probe allowing (F) Raman mapping with perfectly matched location relative to the qBEi image (examples shown).

The Renishaw inLux interface was connected to a Virsa^TM^ Raman spectrometer (Renishaw plc) by optical fibres. The spectrometer was equipped with 785 nm single point laser excitation and a 1500 l/mm diffraction grating. In this configuration, the laser spot size (spatial resolution) was < 2 µm. Briefly, a ROI consisting of 15 by 15 pixels was drawn on the sample surface and analysed first by qBEI.Raman spectra were then collected with exposure to 20 mW 785 nm laser excitation, and 5 seconds integration within the spectral range of 350-1800 cm^-1^. Two step sizes were used during Raman mapping: 2 µm, leading to a 30 µm x 30 µm map, and 9 µm, corresponding to a 135 µm x 135 µm map. These values are inherently related to the resolution of the resulting Raman image, with smaller step sizes producing images with higher resolution, *i.e*., more points per area mapped.

### 2.4. Raman post-processing

Raman maps were post-processed in a lab-made script using Matlab 2023b. Briefly, individual spectra were corrected to remove cosmic rays using the v_1_PO4 band as reference and were baseline corrected using a non-quadratic cost function ^23^. Bone spectra were smoothed using the Savitzky-Golay algorithm (span length 21, degree 4). As pMMA Raman spectrum exhibited significant emission in the spectral range used to assess the organic phase of the bone matrix, resin contribution was subtracted for all spectra using the ∼812 cm^-1^ peak. Table 1 describes the following peak intensity ratios computed.

**Table 1:** Peak intensity ratios used in the present studies. Pro: Proline, Hyp: Hydroxyproline, Phe: Phenylalanine.

| Parameter name | Wavenumber (cm<sup>-1</sup>) |
| --- | --- |
| $\nu_2\text{PO}_4/\text{Amide III}$ | ~428 / ~1240 |
| $\nu_1\text{PO}_4/(\text{Pro}+\text{Hyp})$ | ~960 / (~854 + ~872) |
| $\nu_1\text{PO}_4/\text{Phe}$ | ~960 / ~1003 |
| $\nu_1\text{PO}_4/\text{Amide III}$ | ~960 / ~1240 |
| $\nu_1\text{PO}_4/\text{CH}_2$ | ~960 / ~1450 |
| $\nu_1\text{PO}_4/\text{Amide I}$ | ~960 / ~1668 |

### 2.5. Statistical analysis

All statistical analyses were performed using GraphPad Prism 8.0. The Pearson correlation coefficient was used to determine the strength of correlation between mineral-to-matrix ratio measured by qBEI or Raman spectroscopy. Linear regression between mineral-to-matrix ratio derived from qBEI and mineral-to-matrix ratios measured by Raman spectroscopy was computed and estimated with a F-test and considered significant at p<0.05.

## 3. RESULTS

As qBEI is not often performed in low vacuum mode, we thought to determine whether BMDD was affected by changes in the SEM chamber vacuum. The same bone specimen was investigated at 2.10^−4^ Pa and 50 Pa with two separate SEM microscopes.

As shown in Table 2, BMDD was very similar between the two vacuum modalities, especially if one takes into account thatthe ROI of the bone specimen that had been evaluated between both vacuum modalities was not per se the same due to polishing of the surface.

**Table 2:** BMDD parameters in high and low vacuum modes. In high and low vacuum mode, the conversion equation of gray level (GL) to wt% Ca was: wt% Ca = 0.164.GL-4.26.

| BMDD parameters | High vacuum mode | Low vacuum mode |
| --- | --- | --- |
| $\text{Ca}_{\text{mean}}$ (wt% Ca) | 24.3 | 24.7 |
| $\text{Ca}_{\text{peak}}$ (wt% Ca) | 25.5 | 25.8 |
| $\text{Ca}_{\text{width}}$ (wt% Ca) | 3.0 | 3.5 |
| $\text{Ca}_{\text{low}}$ (wt% Ca) | 2.2 | 3.8 |
| $\text{Ca}_{\text{high}}$ (wt% Ca) | 40.1 | 61.4 |

Since the low vacuum SEM was also fitted with the Raman interface, we thought to determine which of the Raman conventional PO_4_/organic matrix parameters correlated best with the calcium content measured by qBEI. Because the degree of matrix mineralization is determined in Raman spectroscopy as the ratio between mineral and organic matrix, we also transformed the calcium content map, obtained by qBEI, to a wt% mineral / wt% organic matrix map. We binned the qBEI image acquired in low vacuum to match the step size of Raman map. Figure 2 shows the mineral/organic ratio obtained from qBEI image and Raman spectroscopy at a step size of 2 µm. A strong linear relationship was observed for all mineral-to-matrix ratios. Table 3 summarizes the Pearson correlation coefficients and evidences a strong association, ranking from v_1_PO_4_/CH_2_ (R = 0.880), v_1_PO_4_/Phe (R = 0.865), v_1_PO_4_/Amide I (R = 0.858), v_1_PO_4_/(Pro+Hyp) (R = 0.852), v_1_PO_4_/Amide III (R = 0.846) and v_2_PO_4_/Amide III (R = 0.690). In order to assess whether the spatial resolution of the Raman mapping analysis was affecting the outcome, we repeated the same Raman analysis but with a step size of 9 µm. Interestingly, mineral-to-matrix ratios were still linearly correlated (p<0.001 for all linear regression) with wt%Mineral / 100-wt%Mineral (Figure 3). At this spatial resolution, Raman and qBEI mineral-to-matrix ratios were still strongly associated (Table 4) but with lower Pearson correlation coefficients, as represented by v_1_PO_4_/CH_2_ (R = 0.563), v_1_PO_4_/Amide I (R = 0.512), v_1_PO_4_/Amide III (R = 0.488), v_1_PO_4_/Phe (R = 0.455), v_2_PO_4_/Amide III (R = 0.426), and v_1_PO_4_/(Pro+Hyp) (R = 0.403).

**Table 3:** Pearson correlation coefficient between qBEI and Raman mineral-to-matrix ratios at 2 µm step size.

|  | Wt% mineral-to-matrix ratio | p value |
| --- | --- | --- |
| $v_1\text{PO}_4/\text{Amide I}$ | 0.858 | <0.001 |
| $v_1\text{PO}_4/\text{CH}_2$ | 0.880 | <0.001 |
| $v_1\text{PO}_4/\text{Amide III}$ | 0.846 | <0.001 |
| $v_1\text{PO}_4/\text{Phe}$ | 0.865 | <0.001 |
| $v_1\text{PO}_4/(\text{Pro}+\text{Hyp})$ | 0.852 | <0.001 |
| $v_2\text{PO}_4/\text{Amide III}$ | 0.690 | <0.001 |

**Table 4:** Pearson correlation coefficient between qBEI and Raman mineral-to-matrix ratios at 9 µm step size.

|  | Wt% mineral-to-matrix ratio | p value |
| --- | --- | --- |
| $v_1\text{PO}_4/\text{Amide I}$ | 0.512 | $<0.001$ |
| $v_1\text{PO}_4/\text{CH}_2$ | 0.563 | $<0.001$ |
| $v_1\text{PO}_4/\text{Amide III}$ | 0.488 | $<0.001$ |
| $v_1\text{PO}_4/\text{Phe}$ | 0.455 | $<0.001$ |
| $v_1\text{PO}_4/(\text{Pro}+\text{Hyp})$ | 0.403 | $<0.001$ |
| $v_2\text{PO}_4/\text{Amide III}$ | 0.426 | $<0.001$ |

**Figure 2:**
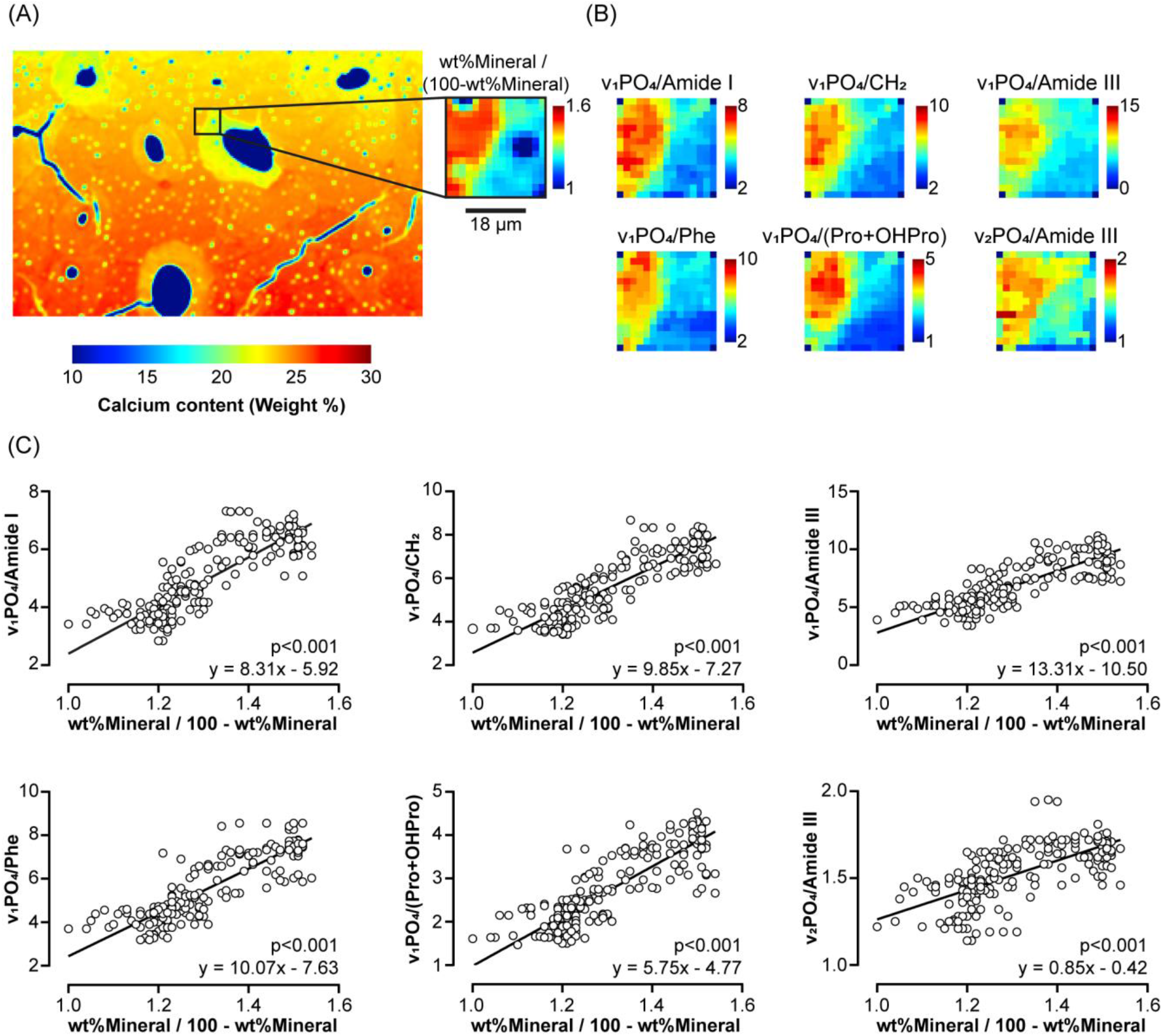
Comparison between mineral/organic ratios obtained by qBEI and Raman spectroscopy at 2 µm step size. (A) the wt% calcium map was obtained from qBEI and a ROI representing a 30 µm x 30 µm area was drawn at the interface between osteon and interstitial bone (black square). Pixels in the ROI were binned to match the spatial resolution of the Raman image and expressed as the ratio between wt% mineral and 100-wt% mineral, representing the mineral-to-matrix ratio. (B) Six mineral-to-matrix ratios were calculated from the Raman image. (C) Regression analyses between the mineral-to-matrix ratio calculated from qBEI and the mineral-to-matrix ratios derived from the Raman map were performed. Linear regression was evaluated by F-test and considered significant at p<0.05.

**Figure 3:**
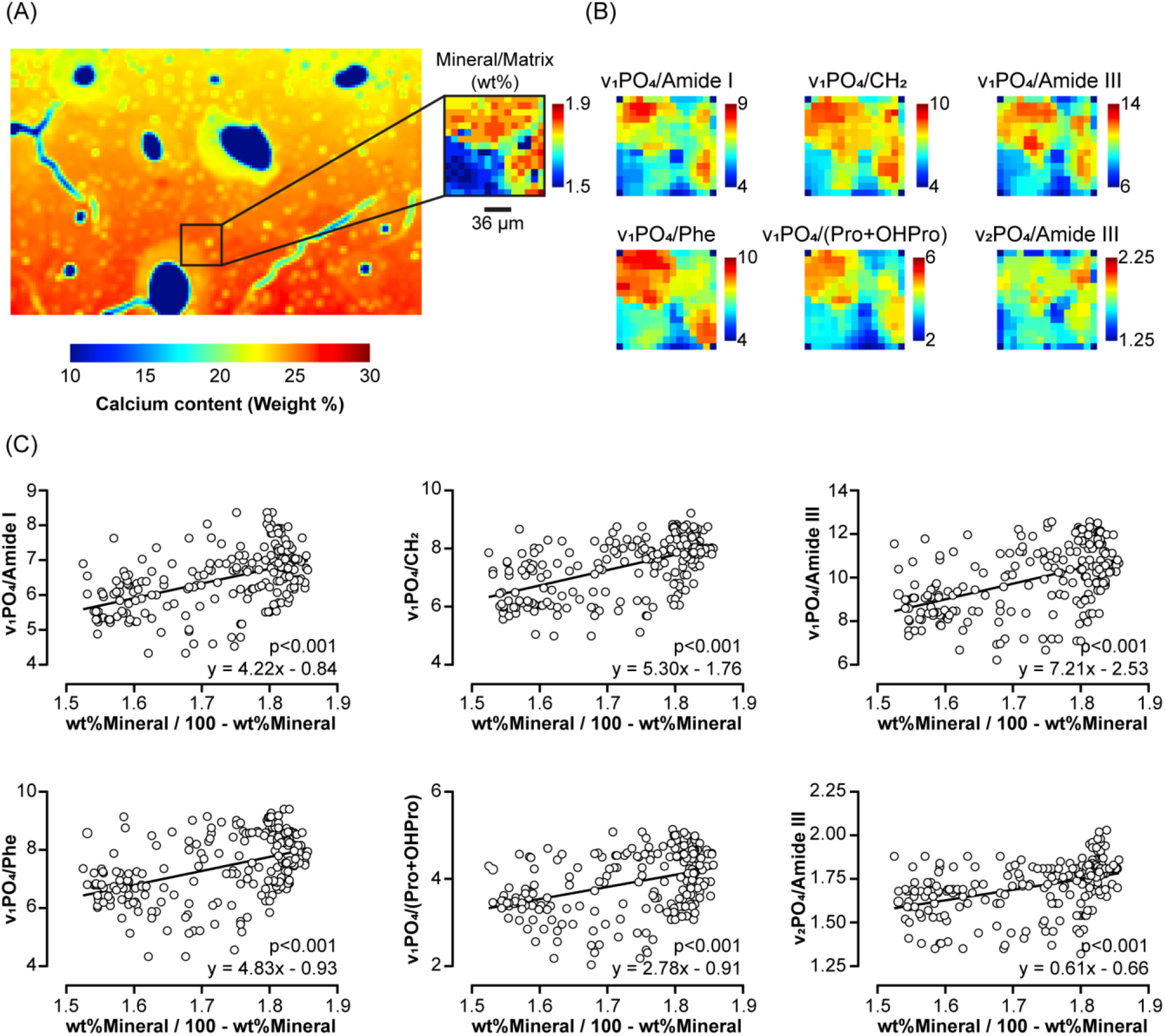
Comparison between mineral/organic ratio obtained by qBEI and Raman spectroscopy at 9 µm step size. (A) the wt% calcium map was obtained from qBEI and a ROI was drawn at the interface between osteon and interstitial bone (black square). Pixels in the ROI were binned to match the spatial resolution of the Raman image (135 µm x 135 µm) and expressed as the ratio between wt% mineral and 100-wt% mineral, representing a mineral-to-matrix ratio. (B) Six mineral-to-matrix ratios were calculated from the Raman image. (C) Regression analyses between the mineral-to-matrix ratio computed from qBEI and the mineral-to-matrix ratios derived from Raman map were performed. Linear regression was evaluated by F-test and considered significant at p<0.05.

## 4. DISCUSSION

Bone resistance to fracture is related to the quantity of bone material, the spatial arrangement of bone packets, and the properties of the bone material itself. The latter has emerged in the last 20 years as an important contributor to bone strength and several studies have linked the mineral-to-matrix ratio to bone strength and toughness ^3-6,24,25^. In the present study, we investigated the correlation between mineral-to-matrix ratio (wt%Mineral / 100- wt%Mineral) estimated by qBEI, and several mineral-to-matrix ratios, using the v_1_PO_4_ or v_2_PO_4_ and Raman peaks assigned to organic moieties. The highest correlation, regardless of the spatial resolution, was encountered for v_1_PO_4_/CH_2_. Previously, this mineral-to-matrix ratio has been positively associated with elastic modulus, yield strength and ultimate strength ^25^. Furthermore, Taylor et al. previously showed a strong association between ash weight, an indicator of mineral content, and the Raman v_1_PO_4_/CH_2_ ratio ^26^. Altogether, this supports the use of v1PO4/CH2 as a good indicator of tissue mineral density.

Intriguingly, v_1_PO_4_/(Pro+Hyp) and v_2_PO_4_/Amide III ratios, have been reported previously to be less sensitive to laser polarization and Raman intensity fluctuations. The v_1_PO_4_/(Pro+Hyp) ratio was also previously shown to strongly correlate with ash weight, tissue mineral density and mineral-to-matrix ratio determined by Fourier-transform infrared spectroscopy (FTIR), as well as elastic modulus, yield strength and ultimate strength ^4,5,12,26,27^. However, in the present study, regardless of spatial resolution, these two ratios exhibited the lowest Pearson correlation coefficients, despite the still positive linear association with wt% mineral-to-matrix ratio determined by qBEI. The reason for the discrepancy between both studies is undetermined at present and we can only make assumptions at this stage. Indeed, an explanation could rely on the fact that the region of interest investigated between qBEI and Raman was accurately matched in the present study. Additionally, this discrepancy could also be explained by the optics of the inLux – Virsa system and the use of optical fibers rather than a conventional microscope objective. Indeed, the fibre optics, as well as the parabolic mirror, used in the inLux system could lead to a scrambling effect on both laser delivery and return fibre to the Ramanspectrophotometer and in turn be less sensitive to polarisation than conventional objective-based Raman system.

Although the utility of the v_2_PO_4_/Amide III has been demonstrated due to minimization of the dependence of Raman intensity to laser polarization ^15^, the spectral position of the v_2_PO_4_ vibrational mode requires longer scan times to capture the low wavenumber regime. In turn, this could be judged disadvantageous for Raman imaging as it would increase the analysis time significantly. Furthermore, the v_1_PO_4_ vibrational mode represents a better alternative that strongly correlates with the degree of mineralization for instruments without polarizing and depolarizing optics, as highlighted in Tables 3 and 4.

Interestingly, the linear correlation between the mineral-to-matrix ratio determined by Raman spectroscopy and qBEI decreased with the reduction in spatial resolution. The typical laser spot size when using a 785 nm laser excitation with the Raman system utilised in this study was estimated at ∼2 µm which in turn dictated the spatial resolution of the Raman image. As the changes in the bone material show a certain heterogeneity, it is likely that binning qBEI pixels, and hence averaging the wt% Ca in the binned pixels, does not totally reflect the heterogeneity of the bone material. A step size higher than the qBEI pixel size implies *de facto* that only a subregion of the pixel has been evaluated and it does not represent the full heterogeneity. Of course, the step size of SEM and Raman images could be lowered down to the nanometer range, however, this would imply an oversampling of the investigated Raman pixels and it would be interesting to assess whether strong correlations could be achieved in this scenario.

The strength of the present study relies on the use of the in-SEM Raman interface that allows Raman spectra to be acquired directly in the SEM chamber and co-localised with the SEM scanned area. One of the limits of performing qBEI and Raman spectroscopy on two different instruments lies in the difficulty of accurately identifying the same ROIs in bone samples. This was demonstrated in the previous seminal paper by Roschger et al., who reported an accuracy of ∼3 µm in identifying and replacing ROIs ^22^. Therefore, the inLux SEM Raman interface is particularly advantageous in (i) reducing variabilities in ROIs as both investigations are performed at the same location, (ii) increasing processing time, as there is no longer the need to remove the specimen from the SEM chamber to bring it to the Raman microscope, and (iii) avoiding contamination and sample damaging while transferring between systems. Indeed, in-SEM Raman spectroscopy has been reported as a superior technique than separated SEM and Raman imaging for correlative microscopy This is of interest as there is increasing demand for the determination of bone material properties and alterations in bone specimens from patients suffering of idiopathic osteoporosis and bone fragility^28^. A limitation of this study is that the native tissue specimens were dehydrated and embedded in polymethylmethacrylate. The pMMA contribution was numerically subtracted based on reference spectra collected next to the bone specimen and the predominant ∼812 cm^-1^ peak of this material. However, pMMA spectrum exhibits several overlapping vibrations with both the v_1_PO_4_ peak and the organic moieties, especially amide I and III, and the CH_2_ peaks. Nevertheless, in the context of clinical diagnostics on bone biopsies, it is likely that the bone specimen would be processed in a similar manner to the one presented here, in which a strong relationship was evidenced between Raman spectroscopy and qBEI for determining the degree of mineralization of the bone matrix.

In conclusion, we have demonstrated in the present study that several mineral-to-matrix ratios, determined with a co-localised Raman system placed inside the SEM chamber, were strongly correlated to the degree of mineralization measured by quantitative backscattered electron imaging, one of the gold standard methodologies to determine tissue mineralization. This is important with respect to the growing demand in bone pathology units to assess alterations of bone material properties and diagnose bone fragility disorders.

## 5 AUTHOR CONTRIBUTION

**Guillaume Mabilleau:** Conceptualization, Investigation, Formal analysis, Writing – Original draft; **Dale Boorman:** Investigation, Formal analysis, Writing – Review & Editing; **Jorge Diniz:** Investigation, Formal analysis, Writing - Review & Editing

## 6 ACKNOWLEDGEMENTS

The authors thank the HiMolA Platform, Inserm UMR_S 1229 RMeS and University of Angers for assistance with bone embedding and preparation. Part of this work was supported by an institutional grant from the University of Angers.

